# Description of *Corynebacterium rouxii* sp. nov., a novel member of the *diphtheriae* species complex

**DOI:** 10.1101/855692

**Authors:** Edgar Badell, Mélanie Hennart, Carla Rodrigues, Virginie Passet, Melody Dazas, Leonardo Panunzi, Valérie Bouchez, Annick Carmi-Leroy, Julie Toubiana, Sylvain Brisse

## Abstract

A group of six clinical isolates previously identified as *Corynebacterium diphtheriae* biovar Belfanti, isolated from human cutaneous or peritoneum infections and from one dog, were characterized by genomic sequencing, biochemical analysis and mass spectrometry (MALDI-TOF MS). The six isolates were negative for the diphtheria toxin gene. Phylogenetic analyses showed that the six isolates (including FRC0190^T^) are clearly demarcated from *C. diphtheriae, C. belfantii, C. ulcerans* and *C. pseudotuberculosis*. The average nucleotide identity of FRC0190^T^ with *C. diphtheriae* NCTC 11397^T^ was 92.6%, and was 91.8% with *C. belfantii* FRC0043^T^. *C. diphtheriae* subsp. *lausannense* strain CHUV2995^T^ appeared to be a later heterotypic synonym of *C. belfantii* (ANI, 99.3%). Phenotyping data revealed an atypical negative or heterogeneous intermediate maltose fermentation reaction for the six isolates. MALDI-TOF MS differentiated the new group from the other *Corynebacterium* taxa by the presence of specific spectral peaks. *rpoB* sequences showed identity to atypical, maltose-negative *C. diphtheriae* biovar Belfanti isolates previously described from two cats in the USA. We propose the name *Corynebacterium rouxii* sp. nov. for the novel group, with FRC0190^T^ (= CIP 111752^T^ = DSM 110354^T^) as type strain.

## Introduction

The genus *Corynebacterium* currently includes approximately 111 species (1–3). The most important human pathogen of the genus is *Corynebacterium diphtheriae*, which causes diphtheria (2, 4). *C. diphtheriae* is genetically heterogeneous (5–8) and four biovars were defined: Gravis, Mitis, Belfanti and Intermedius (9–11), the latter being almost never reported in recent literature. In 2010, maltose-non fermenting strains of *C. diphtheriae* biovar Belfanti were reported from two cats in the USA, and were shown to have a divergent *rpoB* sequence (12). In 2018, some biovar Belfanti isolates were classified as a novel species, *C. belfantii* (3), with 94.85% average nucleotide identity (ANI) with *C. diphtheriae*. Slightly later, *C. diphtheriae* subsp. *lausannense* was also proposed for strains of biovar Belfantii (13). The *tox* gene, which codes for diphtheria toxin, is carried on a corynephage that can lysogenize strains of *C. diphtheriae*. However, the *tox* gene was rarely reported in isolates of biovar Belfanti (5, 14, 15) and no strain of *C. belfantii* or *C. diphtheriae* subsp. *lausannense* was described as *tox*-positive (3)(12). The *tox* gene can also be harboured by strains of *C. ulcerans* and *C. pseudotuberculosis*, two species that are phylogenetically close to *C. diphtheriae* and *C. belfantii* (16). Together, the above-mentioned species constitute a single phylogenetic clade nested within the *Corynebacterium* genus. We refer to this clade as the *C. diphtheriae* complex.

Here, we define the taxonomic status of six isolates initially identified as *C. diphtheriae* biovar Belfanti, isolated from five human infections and one dog in France.

## Material and Methods

We compared the six atypical clinical isolates, among which strain FRC0190^T^, with 13 *C. diphtheriae* strains of biovars Gravis or Mitis (including *C. diphtheriae* type strain NCTC 11397^T^) and 8 strains previously (3) identified as *C. belfantii* (including the type strain FRC0043^T^; **Table 1; Table S1**). Type strains of *C. ulcerans* (CIP 106504^T^ = NCTC 7910^T^) and of *C. pseudotuberculosis* (CIP 102968^T^ = ATCC 19410^T^) were also included for comparison.

**Table 1:** Strains used in this study and their characteristics.

| Table 1 : Strains used in this study and their characteristics |  |  |  |  |  |  |  |  |  |  |
| --- | --- | --- | --- | --- | --- | --- | --- | --- | --- | --- |
| Isolate & | Species | biovar # | Isolation year | Country | Geographic origin @ | tox gene | Isolation source | Disease | Polymicrobial infection | Reference |
| FRC0190<sup>T</sup> | *C. rouxii* | Belfanti | 2013 | France | Lot, Cahors | Negative | Cutaneous | Foot ulceration, chronic arteritis |  | This study |
| FRC0071 | *C. rouxii* | Belfanti | 2011 | France | Haute-Garonne, Toulouse | Negative | Cutaneous | Leg ulceration on chronic arteritis - diabetes |  | This study |
| FRC0284 | *C. rouxii* | Belfanti | 2015 | France | Rhone, Lyon | Negative | Cutaneous | Limb amputation - vasculitis |  | This study |
| FRC0297 | *C. rouxii* | Belfanti | 2015 | France | Herault, Beziers | Negative | Ascitic fluid | Spontaneous peritonitis |  | This study |
| FRC0412 | *C. rouxii* | Belfanti | 2016 | France | Lot, Cahors | Negative | Cutaneous | Purulent orbital cellulitis (dog) |  | This study |
| FRC0527 | *C. rouxii* | Belfanti | 2017 | France | Savoie, Chambéry | Negative | Cutaneous | Foot ulceration on chronic arteritis |  | This study |
| FRC0043<sup>T</sup> | *C. belfantii* | Belfanti | 2009 | France | Corrèze, Brives | Negative | Pharyngeal membrane | Laryngitis |  | Dazas *et al.* 2018 IJSEM |
| 06-4305 | *C. belfantii* | Belfanti | 2006 | France | Rhone, Lyon | Negative | Expectoration | Bronchopathy |  | Dazas *et al.* 2018 IJSEM |
| 00-0744 | *C. belfantii* | Belfanti | 2000 | France | Calvados, Caen | Negative | Expectoration | Cystic fibrosis |  | Dazas *et al.* 2018 IJSEM |
| FRC0074 | *C. belfantii* | Belfanti | 2011 | France | Cote d'Or, Dijon | Negative | Expectoration | Cystic fibrosis |  | Dazas *et al.* 2018 IJSEM |
| FRC0223 | *C. belfantii* | Belfanti | 2014 | France | Pas-de-Calais, Coquelles | Negative | Sinusal swab | Sinusitis |  | Dazas *et al.* 2018 IJSEM |
| 05-3187 | *C. belfantii* | Belfanti | 2005 | France | Seine-Maritime, Rouen | Negative | Nasal swab | Rhinitis |  | Dazas *et al.* 2018 IJSEM |
| FRC0250 | *C. belfantii* | Belfanti | 2014 | France | Bas-Rhin, Strasbourg | Negative | Bronchoalveolar wash | Pneumonia |  | Dazas *et al.* 2018 IJSEM |
| FRC0301 | *C. belfantii* | Belfanti | 2015 | France | Calvados, Lisieux | Negative | Expectoration | n.a. |  | Dazas *et al.* 2018 IJSEM |
| NCTC 11397<sup>T</sup> | *C. diphtheriae* | Gravis | 1969 | USA | New York, USA | Negative | n.a. | n.a. |  | Dazas *et al.* 2018 IJSEM |
| NCTC 13129 | *C. diphtheriae* | Gravis | 1997 | United Kingdom | Unknown | Positive | Pharyngeal membrane | Diphtheria |  | Dazas *et al.* 2018 IJSEM |
| FRC0336 | *C. diphtheriae* | Gravis | 2015 | France | Ille-et-Vilaine, Rennes | Positive | Cutaneous | Leishmaniasis | *Leishmania spp.* | Dazas *et al.* 2018 IJSEM |
| FRC0304 | *C. diphtheriae* | Gravis | 2015 | France | La Reunion, St Denis | Negative | Cutaneous | Bullous skin lesion |  | Dazas *et al.* 2018 IJSEM |
| FRC0375 | *C. diphtheriae* | Mitis | 2015 | France | Oise, Creil | Positive | Cutaneous | Ankle ulceration |  | Dazas *et al.* 2018 IJSEM |
| FRC0432 | *C. diphtheriae* | Mitis | 2016 | France | Seine-et-Marne, Vaires sur Marne | Negative | Cutaneous | Purulent scalp skin injury |  | Dazas *et al.* 2018 IJSEM |
| FRC0157 | *C. diphtheriae* | Mitis | 2013 | France | Paris | Negative | Cutaneous | Left ankle wound | *S. pyogenes* | Dazas *et al.* 2018 IJSEM |
| FRC0132 | *C. diphtheriae* | Mitis | 2012 | France | Yvelines, Le Chesnay (return from Mali) | Negative | Cutaneous | Necrotic lesions |  | Dazas *et al.* 2018 IJSEM |
| FRC0036 | *C. diphtheriae* | Mitis | 2009 | France | Mayotte, Mamoudzou | Negative | Cutaneous | Burn wound |  | Dazas *et al.* 2018 IJSEM |
| FRC0154 | *C. diphtheriae* | Mitis | 2012 | France | Haut-Rhin, Colmar | Positive | Cutaneous | Cutaneous infection | *S. pyogenes* | Dazas *et al.* 2018 IJSEM |
| FRC0049 | *C. diphtheriae* | Mitis | 2009 | France | Mayotte, Mamoudzou | Positive | Cutaneous | Genital lesion |  | Dazas *et al.* 2018 IJSEM |
| FRC0430 | *C. diphtheriae* | Mitis | 2016 | France | Rhone, Bron | Positive | Cutaneous | Leg ulcerations | *S. pyogenes* | Dazas *et al.* 2018 IJSEM |
| FRC0436 | *C. diphtheriae* | Mitis | 2016 | France | Ille-et-Vilaine, Rennes | Positive | Cutaneous | Cutaneous infection | *S. pyogenes* | Dazas *et al.* 2018 IJSEM |
| ATCC 19410<sup>T</sup> | *C. pseudotuberculosis* | not applicable | 1931 | n.a. | South America | Negative | Infected gland (sheep) | n.a. |  | PMID: 13882624 (Cummins, 1962) |
| NCTC 7910<sup>T</sup> | *C. ulcerans* | not applicable | 1948 | United Kingdom | n.a. | Negative | Throat | n.a. |  | PMID: 7729671 (Riegel et al., 1995) |
| # Biovar of *C. diphtheriae* as classically defined |  |  |  |  |  |  |  |  |  |  |
| & FRC: collection of the French National Reference Center for the Corynebacteria of the *C. diphtheriae* complex; NCTC: National Collection of Type Cultures (Public Health England); ATCC: American Type Culture Collection |  |  |  |  |  |  |  |  |  |  |
| @ Geographic origin for French isolates is given as "French Department, city" |  |  |  |  |  |  |  |  |  |  |
| \$ Sequence type is defined at https://pubmlst.org/cdiphtheriae/ | | | | | | | | | | |
| n.a.: not available |  |  |  |  |  |  |  |  |  |  |
| NA: not applicable |  |  |  |  |  |  |  |  |  |  |

Clinical samples or isolates were sent to the French National Reference Centre for Corynebacteria of the *diphtheriae* complex for isolation and/or characterization, respectively. Oxoid’s Tinsdale agar with supplement medium (Thermo Fisher Diagnostics, Dardilly, France) was used to isolate *C. diphtheriae* from clinical samples. Isolates were frozen in Brain-Heart-Infusion (BHI) medium containing 30% of glycerol and stored at -80°C prior to this study. After thawing, isolates were grown at 37°C on tryptose-casein soy agar plates during 24 hours. DNA was extracted from a few colonies with the DNA Blood and Tissue kit (Qiagen, Hilden, Germany). The six isolates were identified as *C. diphtheriae* or *C. belfantii* by multiplex polymerase chain reaction (PCR) combining a *dtxR* gene fragment specific for *C. diphtheriae* (15) and a multiplex PCR (17, 18) that targets a fragment of the *pld* gene specific for *C. pseudotuberculosis*, the gene *rpoB* (amplified in all species of the *C. diphtheriae* complex) and a fragment of 16S rRNA gene specific for *C. pseudotuberculosis* and *C. ulcerans*. The *tox* gene was also detected by PCR (19). These PCR results were confirmed using a more recent four-plex qPCR (20).

For biochemical identification, standard methods were used (20). More specifically, strains were characterized for pyrazinamidase, urease, nitrate reductase and for utilization of maltose and trehalose using API coryne strips (BioMérieux, Marcy l’Etoile, France) and the Rosco Diagnostica reagents (Eurobio, Les Ulis, France). The Hiss serum water test was used for glycogen fermentation. The biovar of isolates was determined based on the combination of nitrate reductase (positive in Mitis and Gravis, negative in Belfanti) and glycogen fermentation (positive in Gravis only).

Antimicrobial susceptibility was characterized by the disk diffusion method using impregnated paper disks (Bio-Rad, Marnes-la-Coquette, France) and minimum inhibitory concentrations were determined using E-test strips (BioMérieux, Marcy l’Etoile, France). The sensitivity was interpreted using CA-SFM/EUCAST V.1.0 (Jan 2019) criteria for *Corynebacterium* (https://www.sfm-microbiologie.org/wp-content/uploads/2019/02/CASFM2019_V1.0.pdf). Susceptibility was tested for the following antimicrobial agents: fosfomycin, vancomycin, kanamycin, gentamycin, penicillin G, oxacillin, amoxicillin, imipenem, cefotaxime, clindamycin, azithromycin, spiramycin, clarithromycin, erythromycin, clindamycin, ciprofloxacin, trimethoprim-sulfamethoxazole, trimethoprim, sulfonamide, pristinamycin, rifampicin and tetracycline.

MALDI-TOF mass spectrometry (MS) was used for identification confirmation. For this purpose, an overnight culture on Trypto-Casein-Soy Agar (TSA) (37°C) was used to prepare the samples accordingly to the ethanol/formic acid extraction procedure proposed by in the manufacturer recommendations (Bruker Daltonics, Bremen, Germany). The cell extracts were then spotted onto an MBT Biotarget 96 target plate, air dried and overlaid with 1 μL of a saturated α-cyano-4-hydroxycinnamic acid (HCCA). 24 mass spectra per strain were acquired on a Microflex LT mass spectrometer (Bruker Daltonics, Bremen, Germany). Re-analysis of the spectra was performed for the purpose of this work. Spectra were first preprocessed by applying smoothing and baseline subtraction with FlexAnalysis software using default parameters, exported as text files from the Brucker system and then imported and analyzed in a dedicated BioNumerics v7.6.3 (Applied-Maths, Belgium) database following the protocol described by Rodrigues *et al.* (22). To allocate proteins to the specific peaks detected, we extracted all the molecular weights from the genomes of the type strains (NTCT11397^T^, FRC0043^T^ and FRC0190^T^) using a Biopython script (https://biopython.org/DIST/docs/api/Bio.SeqUtils-module.html) and performed sequence alignments with ClustalW for the candidate proteins.

Genomic sequencing was performed from Nextera XT libraries using a NextSeq-500 instrument (Illumina, San Diego, USA) with a 2 × 150 nt paired-end protocol. Contig sequences were assembled using SPAdes v3.12.0 (23) (**Table S1**). JSpeciesWS (24) was used to calculate the BLAST-based average nucleotide identity (ANIb). BLAST was used to extract 16S rRNA and *rpoB* sequences from genome assemblies and to determine the presence or absence of the *narIJHGK* nitrate reduction gene cluster using as query the cluster of strain NCTC 13129 (RefSeq accession number: DIP_RS13820 to DIP_RS13845) (25). *rpoB* and 16S rRNA gene sequences of atypical *C. belfantii* strains from cats (12) were included for comparison. For genome-based phylogenetic analysis, the pairwise *p*-distance (*i.e.*, proportion of aligned nucleotide differences) between each pair of genomes was estimated based on Mash (26) using a multiple hit correction (27) with JolyTree (https://gitlab.pasteur.fr/GIPhy/JolyTree). For 16S rRNA and rpoB gene sequences, sequences were aligned with MAFFT v7.407 (28) and the resulting alignment was used for phylogenetic tree inference with IQ-TREE v1.6.7.2 (29) using the GTR+I+G4 model.

### Data accessibility

Sequence data generated in this study were deposited in the European Nucleotide Archive database and are accessible under project number PRJEB22103. The EMBL (GenBank/DDBJ) accession numbers of the genomic sequences released in this study are ERS3795539 to ERS3795544. The annotated genomic sequence of strain FRC0190^T^ was deposited in the European Nucleotide Archive and is available under accession number ERZ1195831. *rpoB* and 16S rRNA gene sequences were also submitted individually under accession numbers MN542347 to MN542352 and MN535982 to MN535987, respectively.

## Results and discussion

Six isolates were isolated from five cutaneous lesions and one ascitic fluid sample (**Table 1**). Strikingly, human cutaneous lesions were all ulcerations due to underlying chronic arteritis. Ascitic fluid was sampled on a patient with a suspicion of spontaneous peritonitis. The dog was investigated in the context of purulent orbital cellulitis.

The six isolates were *tox* negative (**Table S1**); more specifically, they were negative for amplification of the expected 910-bp PCR product encompassing fragments A and B of the toxin gene (20) and also negative for the amplification of a 117-bp region of diphtheria toxin fragment A (30) by multiplex qPCR as described in (19). We also confirmed by BLASTN that the *tox* gene sequence (query: *tox* gene sequence from strain NCTC 13129, RefSeq accession number: DIP_RS12515) was absent from the genomic assemblies. After species identification by multiplex PCR, the isolates were positive for *dtxR* and *rpoB* and negative for *C. ulcerans/C. pseudotuberculosis* 16S rDNA and *pld*, leading to initial identification as *C. diphtheriae*. Concordant with this identification, the six isolates were pyrazinamidase, urease and trehalose negative. Upon biotyping, the isolates were nitrate and glycogen negative, a pattern that corresponds to biovar Belfanti. Consistently, the *narKGHIJ* nitrate reduction gene cluster was not detected from the genomic assemblies of these isolates and those of *C. belfantii* (**Table S2**). The phenotypic aspect of colonies on Tinsdale or blood agar medium was undistinctive from *C. diphtheriae* Mitis and Gravis and *C. belfantii*. Interestingly, the maltose test was negative for the six isolates using API coryne (**Table S1**). The same test was atypical using the Rosco Diagnostic method: results showed heterogeneous coloration that was neither as yellow as the typically positive strains, nor as purple as the negative strains (**Figure S3**). This atypical maltose result was not observed using API coryne strips, with which the maltose test was clearly negative for the six isolates.

Regarding their antimicrobial susceptibility, the six isolates were resistant to fosfomycin, as is typical of *Corynebacteria* (31), and were susceptible to all other tested antimicrobial agents with the following exceptions: FRC0284 and FRC0527 were resistant to penicillin (minimum inhibitory concentration: 0.19 mg/L), and FRC0412 was resistant to penicillin and cefotaxime (0.19 mg/L and 1.0 mg/L, respectively).

Genomic sequencing results showed that the six isolates had a genome size of 2.4 Mb on average (**Table S1**), similar to *C. diphtheriae* biovars Mitis and Gravis isolates (average size: 2.45 Mb), but smaller than *C. belfantii* (average size: 2.7 Mb). A genome sequence-based phylogenetic tree (**Figure 1**) revealed three main clades. The first one contained all *C. diphtheriae* Mitis and Gravis isolates, whereas the second comprised all *C. belfantii* isolates, and the third comprised the six maltose-atypical isolates. The mean ANIb value of atypical isolates was 92.4% with the *C. diphtheriae* clade and was 91.4% with *C. belfantii* (**Table 2**). These data indicate that the six isolates forming the atypical clade correspond to a distinct genomic cluster, separated by a level of nucleotide divergence that is well above the currently accepted genomic species threshold of ∼94-96% (32, 33). The atypical clade was genetically homogeneous, with ANIb values among the six isolates ranging from 99.21% to 99.94% (**Table 2**). Phylogenetic analysis of *rpoB* and 16S rRNA coding sequences was consistent with the distinction of the atypical isolates from *C. diphtheriae* and *C. belfantii* (**Figures S1 and S2**). We noted that *rpoB* and 16S rRNA sequences of previously reported atypical biovar Belfanti isolates from cats in the USA (12) were indistinguishable from those of the atypical isolates from France, suggesting that the cat isolates from the USA belong to the same novel group. Supporting this observation, the USA cat isolates were also reported as maltose negative.

**Table 2.** Average nucleotide identity values.

| *Corynebacterium* species | Strain identifier (1) | FRC0071 | FRC0190T | FRC0284 | FRC0297 | FRC0412 | FRC0527 | NCTC11397T | FRC0043T | NCTC7910T | ATCC19410T |
| --- | --- | --- | --- | --- | --- | --- | --- | --- | --- | --- | --- |
| *C. rouxii* sp. nov. | FRC0071 | 100 | 99.60 | 99.89 | 99.21 | 99.37 | 99.94 | 92.30 | 91.22 | 71.34 | 70.94 |
| *C. rouxii* sp. nov. | FRC0190 | 99.68 | 100 | 99.75 | 99.24 | 99.34 | 99.68 | 92.41 | 91.36 | 71.40 | 70.93 |
| *C. rouxii* sp. nov. | FRC0284 | 99.94 | 99.71 | 100 | 99.22 | 99.35 | 99.92 | 92.30 | 91.28 | 71.26 | 70.85 |
| *C. rouxii* sp. nov. | FRC0297 | 99.26 | 99.28 | 99.27 | 100 | 99.29 | 99.26 | 92.44 | 91.43 | 71.16 | 70.67 |
| *C. rouxii* sp. nov. | FRC0412 | 99.45 | 99.26 | 99.40 | 99.23 | 100 | 99.44 | 92.39 | 91.37 | 71.04 | 70.79 |
| *C. rouxii* sp. nov. | FRC0527 | 99.95 | 99.55 | 99.85 | 99.21 | 99.40 | 100 | 92.30 | 91.27 | 71.27 | 70.99 |
| *C. diphtheriae* | NCTC11397T | 92.33 | 92.45 | 92.30 | 92.32 | 92.34 | 92.32 | 100 | 95.07 | 71.29 | 70.86 |
| *C. belfantii* | FRC0043T | 90.92 | 91.06 | 90.93 | 90.99 | 90.99 | 90.92 | 94.77 | 100 | 71.12 | 70.76 |
| *C. ulcerans* | NCTC7910T | 71.42 | 71.51 | 71.43 | 71.32 | 71.30 | 71.42 | 71.40 | 71.31 | 100 | 84.33 |
| *C. pseudotuberculosis* | ATCC19410T | 71.31 | 71.30 | 71.31 | 71.25 | 71.25 | 71.31 | 71.19 | 71.06 | 84.29 | 100 |

(1) A trailing T after the strain identifier denotes that the strain is the type strain of its corresponding taxon

**Figure 1.**
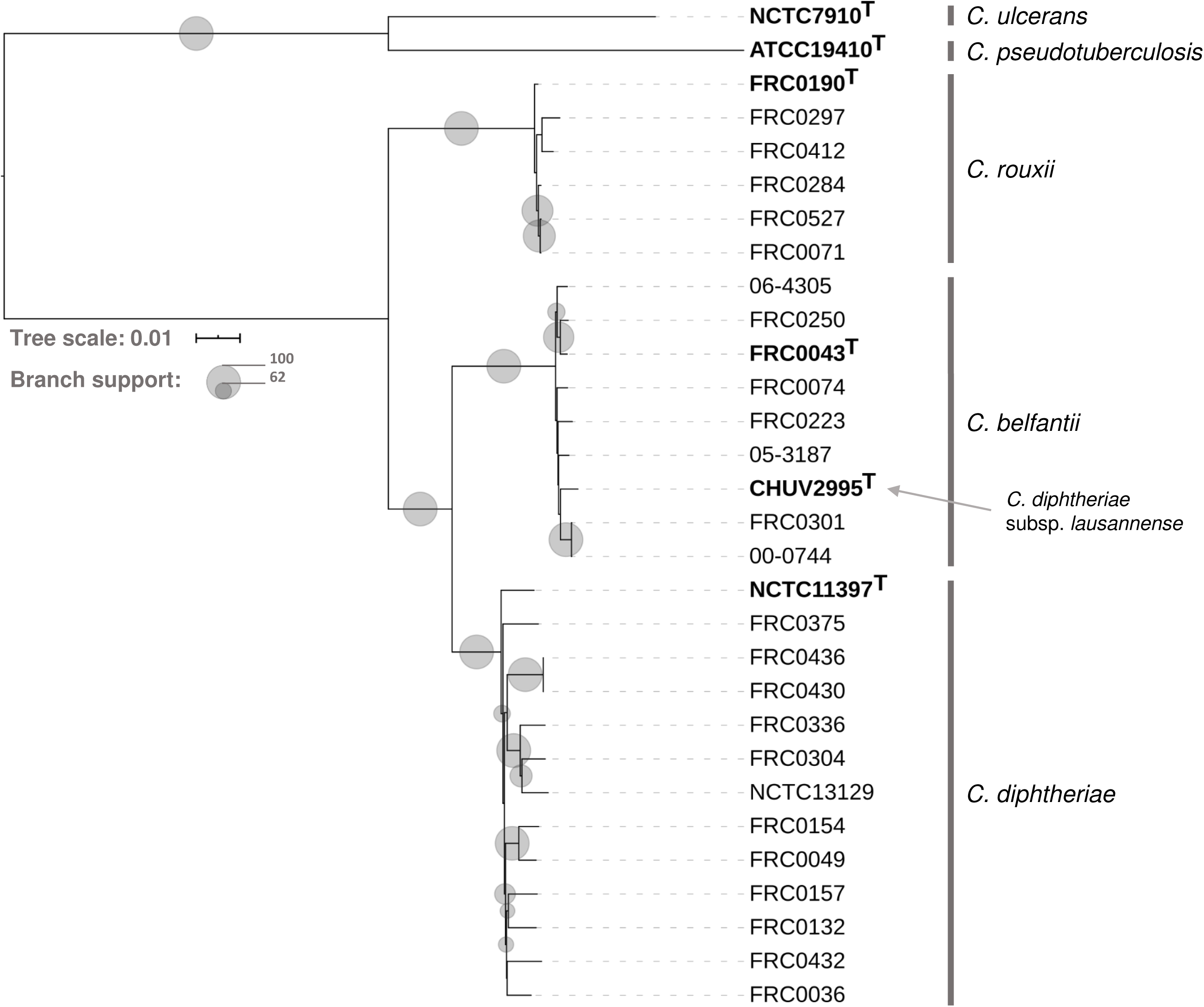
Phylogenetic relationships derived from the analysis of genomic sequences. The phylogenetic tree and branch supports were inferred using JolyTree (27) (https://gitlab.pasteur.fr/GIPhy/JolyTree). Strains *C. ulcerans* NCTC 7910^T^ and *C. pseudotuberculosis* ATCC 19410^T^ were used as outgroup as they are the closest phylogenetic neighbors to *C. diphtheriae, C. belfantii* and *C. rouxii.* Branch support is indicated using grey circles (see key; only values >50 are shown). Each taxonomic type strain is shown in bold; note that *C. diphtheriae* subsp. *lausannense* type strain falls within *C. belfantii*. The scale bar corresponds to an estimated evolutionary distance of 0.01.

Recently, it was proposed that the *C. diphtheriae* taxon should be subdivided into two subspecies, *C. diphtheriae* subsp. *diphtheriae* and *C. diphtheriae* subsp. *lausannense* (13). Here, we observed that the ANI value between the type strains of *C. diphtheriae* subsp. *lausannense* and *C. belfantii* was 99.3%. Besides, the former was positioned within the phylogenetic branch of *C. belfantii* (**Figure 1, S1 and S2**), and the descriptions of both taxa are very similar (3, 13). Given that *C. belfantii* was validly published in October 2018, a few months before the taxonomic proposal *C. diphtheriae* subsp. *lausannense* was validated (https://www.microbiologyresearch.org/content/journal/ijsem/10.1099/ijsem.0.003174; January 2019), the latter subspecies appears to be a later heterotypic synonym of *C. belfantii*.

Based on the MALDI Biotyper Compass database version 4.1.90 (Bruker Daltonics, Bremen, Germany), the six isolates were identified as *C. diphtheriae*. However, detailed analysis of their spectra led to the identification of six pairs of biomarkers (12 peaks corresponding to the same proteins, with either single and double-charged ion forms) corresponding to three different proteins within the range 3255–9495 *m/z*, which were associated either with the group of six isolates or with *C. diphtheriae* and *C. belfantii* (**Figure S4, Table S3**). We presumptively identified the specific biomarkers as two ribosomal proteins, L30 and S20, and one putative stress response protein (CsbD). Consistently, their amino-acid sequences differed between the *C. rouxii* on the one hand, and *C. diphtheriae*/*C. belfantii* on the other hand (**Figure S5**). Based on the current dataset, the specificity and sensitivity of peak distribution among the three species ranged between 95– 100% and 76–100%, respectively (**Table S3**). MALDI-TOF MS thus allows the discrimination between *C. rouxii* and *C. diphtheriae*/*C. belfantii*. These results warrant future updates of reference MALDI-TOF MS databases to incorporate the novel taxon.

Based on the above results, the isolates of the novel clade represent a novel species, which we propose to name *Corynebacterium rouxii.*

### Description of *Corynebacterium rouxii* sp. nov

(rou’.xi.i. N.L. gen. n. *rouxii*, of Roux, a French scientist and former director of Institut Pasteur who made critical contributions to diphtheria toxin discovery and antitoxin treatment).

*C. rouxii* conforms biochemically to the description of *C. diphtheriae* strains belonging to biovar Belfantii (2, 21), except that strains are negative for maltose fermentation (API coryne), being nearly negative or weakly positive with the Rosco Diagnostica test. Key characteristics that distinguish *C. rouxii* from other members of the *C. diphtheriae* complex are specific MALDI-TOF MS biomarkers as described herein. The G+C content of *C. rouxii* genomes ranges from 53.2% to 53.3%, with a value of 53.3% for the type strain. So far, strains were isolated from 5 humans and a dog in France, as well as from two related cats in the USA.

The type strain is FRC0190^T^ (= CIP 111752^T^ = DSM 110354^T^), isolated in 2013 from a foot ulceration reported in Cahors, France. The genome accession number of strain FRC0190^T^ is ERS3795540.

## Acknowledgments

We thank the “Plateforme de Microbiologie Mutualisée” from Institut Pasteur for genomic sequencing, and Prof. Alain Le Coustumier (Centre hospitalier de Cahors, France) for sending the original culture of strain FRC0190^T^.

## Funding information

This work was supported financially by Institut Pasteur and Santé Publique France (Saint-Maurice, France).

## Conflicts of interest

The authors declare that there is no conflict of interest.

## Ethical statement

There is no ethical issue associated with this manuscript.

### Abbreviations

ANI: average nucleotide identity;
MALDI-TOF MS: matrix-assisted laser desorption/ionisation time-of-flight mass spectrometry

---

**Figure S1.** Phylogenetic analysis of 16S rRNA gene sequences.

16S rRNA gene sequences of *C. rouxii* isolates have accession numbers MN535982 to MN535987 (**Table S1**). *C. diphtheriae* strains isolated from domestic cats in the USA (12) were added for comparison (strain codes CD443-CD450; GenBank accession numbers: FJ409572 to FJ409575). Each taxonomic type strain is shown in bold. The scale bar indicates the number of substitutions per site.

**Figure S2.** Phylogenetic analysis of *rpoB* coding sequences.

*rpoB* gene sequences of *C. rouxii* isolates have accession numbers MN542347 to MN542352 (**Table S1**). *C. diphtheriae* strains isolated from domestic cats in the USA (12) were added for comparison (strain codes CD443 and CD450; GenBank accession numbers: FJ415317 and FJ415318). Each taxonomic type strain is shown in bold. The scale bar indicates the number of substitutions per site.

**Figure S3.** Maltose fermentation results.

The maltose test was performed using the Rosco Diagnostica reagents. Tube 1: NCTC 10648, *C. diphtheriae* biovar Gravis; Tubes 2 to 7: *C. rouxii* isolates FRC0071, FRC0297, FRC0284, FRC0527, FRC0190, FRC0412; Tube 8: NCTC 10356, *C. belfantii*; Tube 9: NCTC 12077 (*C. ulcerans*, positive control); Tube 10: NCTC 764 (*C. striatum*, negative control).

**Figure S4**. Peak positions (*m/z*) observed for strains of C. *diphtheriae, C. belfantii* and *C. rouxii*. Stars denote those peaks that are useful for species identification, as detailed in the corresponding supplementary Table.

**Figure S5.** Amino acid sequence alignments and the respective molecular weight of the proteins presumptively associated with specific MALDI-TOF MS peaks detected in *Corynebacterium diphtheriae, C. belfantii* and *C. rouxii*

